# Regulation of sleep plasticity by a thermo-sensitive circuit in *Drosophila*

**DOI:** 10.1101/082487

**Authors:** Angelique Lamaze, Arzu Öztürk-Çolak, Robin Fischer, Nicolai Peschel, Kyunghee Koh, James E.C. Jepson

## Abstract

Sleep is a highly conserved and essential behaviour in many species, including the fruit fly Drosophila melanogaster. In the wild, sensory signalling encoding environmental information must be integrated with sleep drive to ensure that sleep is not initiated during detrimental conditions. However, the molecular and circuit mechanisms by which sleep timing is modulated by the environment are unclear. Here we introduce a novel behavioural paradigm to study this issue. We show that in male fruit flies, onset of the daytime siesta is delayed by ambient temperatures above 29°C. We term this effect Prolonged Morning Wakefulness (PMW). We show that signalling through the TrpA1 thermo-sensor is required for PMW, and that TrpA1 specifically impacts siesta onset, but not night sleep onset, in response to elevated temperatures. We identify two critical TrpA1-expressing circuits and show that both contact DN1p clock neurons, the output of which is also required for PMW. Finally, we identify the circadian blue-light photoreceptor CRYPTOCHROME as a molecular regulator of PMW. We propose a model in which the Drosophila nervous system integrates information encoding temperature, light, and time to dynamically control when sleep is initiated. Our results provide a platform to investigate how environmental inputs co-ordinately regulate sleep plasticity.

## Introduction

Sleep is controlled by circadian and homeostatic mechanisms^1^, and is observed in organisms as divergent as humans and insects^2^. Such a deep conservation suggests a fundamental requirement for sleep in maintaining organismal fitness. Indeed, recent work has demonstrated a key role for sleep in regulating several aspects of nervous system function in vertebrates and invertebrates, including synaptic plasticity, neuronal development and metabolite clearance^3-6^.

The quiescent and high arousal-threshold state that is the primary characteristic of sleep has obvious fitness costs in an environment containing predators and a limited supply of food and mates. Thus, the timing of sleep must be tightly regulated by an array of sensory modalities to match sleep onset to current external conditions. In other words, sleep must be a plastic phenotype that is sensitive to environmental alterations detected by sensory processes. However, the mechanisms by which ethologically-relevant environmental cues modulate sleep timing are unclear. We have used the fruit fly, *Drosophila melanogaster*, to address this issue.

*Drosophila* has emerged as an important model system for investigating genes and circuits that influence the levels, timing, and homeostatic regulation of sleep^7-14^, as well as its neurobiological functions^3-5,15^. *Drosophila* exhibit a daytime siesta and long bouts of consolidated sleep during the night. Components of this sleep pattern are sexually dimorphic, with female flies exhibiting reduced siesta sleep relative to males^7^. As poikilotherms, *Drosophila* physiology is temperature-sensitive, and fruit flies possess an array of thermo-sensory signalling pathways that facilitate adaptive behavioral responses to the surrounding ambient temperature, enabling *Drosophila* to sense attractive or noxious temperatures and initiate appropriate locomotor programs^16,17^. Recent work has also shown that increasing ambient temperature modifies sleep architecture^18-20^. However, the thermo-sensory molecules and circuits that transmit temperature information to sleep regulatory centers remain unclear.

We performed an in-depth analysis of how sleep in *Drosophila* is modified by changes in ambient temperature. Interestingly, we found that the male siesta exhibits a complex response to temperature increases, with the onset advanced during mild increases and delayed by further thermal increases (≥ 30°C). We term this delay in siesta onset due to elevated temperature Prolonged Morning Wakefulness (PMW). Through mutational analysis and circuit mapping approaches, we identify the thermo-receptor, thermo-sensory circuits and downstream sleep-regulatory neurons involved in PMW. We show that temperature increases are encoded by two distinct sub-circuits expressing the TrpA1 thermo-sensor, both of which contact DN1p clock neurons. These, in turn, promote wakefulness during hot mornings. We also demonstrate that loss of the circadian photoreceptor CRYPTOCHROME suppresses PMW. Our work suggests that the *Drosophila* nervous system integrates light, temperature, and temporal information to drive changes in daytime sleep architecture.

## Results

### Increased ambient temperature alters sleep architecture in *Drosophila*

To study how sleep architecture dynamically adapts to environmental changes, we measured sleep levels and timing in adult male flies during two consecutive days. On Day 1 of the experimental paradigm, flies were housed at 22°C, in 12 h light :12 h dark (LD) conditions. On Day 2, flies were exposed to a range of warm temperatures (27-31°C) for 24 h, beginning at Zeitgeber Time 0 (ZT0, lights-on). In these and all subsequent experiments, sleep was defined as 5 min of inactivity, as measured by the *Drosophila* Activity Monitoring (DAM) system, a well-described standard in the field^21^.

Temperature increases resulted in complex changes to the architecture of siesta sleep. Shifting male flies from 22°C to either 27°C or 29°C prolonged siesta sleep towards lights-off (ZT12), yielding a net increase in siesta sleep at 27°C and 29°C compared to 22°C (Fig. 1a, b, f). This may be due to a delay in the initiation of locomotor increases before lights-off (evening anticipation; Fig. S1), an output of the circadian clock previously shown to be temperature-sensitive^22,23^. At 27°C, siesta onset was slightly advanced relative to 22°C (Fig. 1a, d;). However, further increases in temperature shifted siesta onset to later time periods. In particular, at ≥ 30°C we observed a robust delay in siesta onset that was not observed at 29°C (Fig. 1b-d), contributing to a net reduction in siesta sleep at ≥ 30°C (Fig. 1f). With respect to nighttime sleep, we found that heightened ambient temperatures induced a delay in night sleep onset (Fig. 1a-c, e), quantified as the change in latency to initiate sleep between cold and hot days (D Latency). In addition, we observed a roughly linear decrease in night sleep levels in response to increasing temperature levels (Fig. 1g).

**Figure 1:**
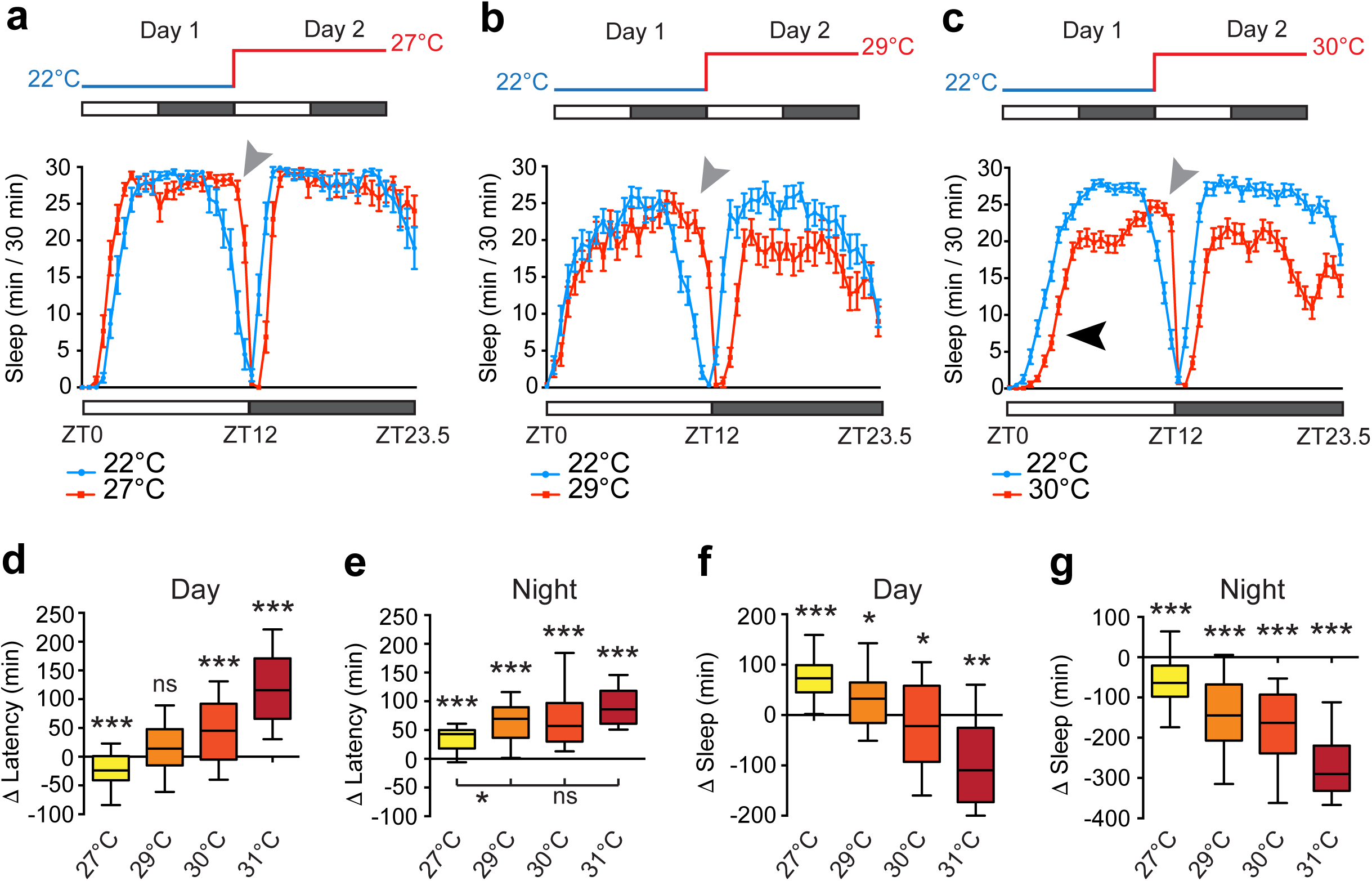
Warm temperatures prolong morning wakefulness in male *Drosophila*. (**a-c**) Average sleep patterns of adult male flies shifted from 22°C to either 27°C (n=20) (**a**), 29°C (**b**) or 30°C (**c**). Sleep traces are presented as mean ± SEM for each time point in these and all subsequent figures. Temperature-shift paradigms are indicated above. Sleep was measured under 12 h light: 12 h dark conditions (white/grey bars) with Zeitgeber Times (ZT) shown below. Black arrowhead indicates the delay of the sleep onset observed at 30°C (PMW). Grey arrowheads indicate the delay of sleep offset induced at 27°C or above. (**d-e**) Change in time taken to the first day sleep episode (**d**) or night sleep episode (**e**) (D Latency) between consecutive 24 h periods at 22°C and 27-31°C. (**f-g**) Difference in total sleep during the day (**f**) or night (**g**) between consecutive 24 h periods at 22°C and 27-31°C. n-values: 27°C, n = 39; 29°C, n = 32; 30°C, n = 79; 31°C, n = 18. In this and all subsequent figures, box plots show the 10^th^, 25^th^, median, 75^th^ and 90^th^ percentiles, and p-values are indicated as follows: *p < 0.05, **p < 0.005, ***p < 0.0005, ns – p > 0.05. Statistical comparisons: (d-g) Wilcoxon signed rank test compared to a theoretical median of zero and (e) Kruskal-Wallis test with Dunn’s post-hoc test.

Thus, temperature increases differentially affect the onset of day versus night sleep, with siesta sleep onset bi-directionally altered by temperature increases within a relatively narrow range, and night sleep onset consistently delayed. To our knowledge, the response of siesta onset to elevated temperatures has not previously been characterized, and for simplicity, we refer to the temperature-induced delay in siesta onset PMW - Prolonged Morning Wakefulness.

To test whether PMW was sex-specific, we also examined sleep in adult female *Drosophila*. At 22°C, female *Drosophila* initiate the siesta later during the day compared to males (Fig. S1), and we did not observe PMW in females when shifted from 22°C to 30°C (Fig. S1). Thus, PMW is sex-specific at 30°C and correlates with a siesta onset occurring earlier in the morning. We also wondered whether PMW represents an acute avoidance response to rapidly increased ambient temperature. To rule out this possibility, we shifted male flies from 22°C to 30°C at ZT12 rather than ZT0 and measured sleep the following morning, after 12 h at elevated temperature (Fig. 2a). Indeed, under these conditions we still observed robust PMW, and the magnitude of PMW was equivalent to that caused by a shift from 22°C to 30°C at ZT0 (Fig. 2b, c). Thus, PMW is not simply a reaction to a rapid environmental change, but is a behavioral response linked to high temperatures during the morning.

**Figure 2:**
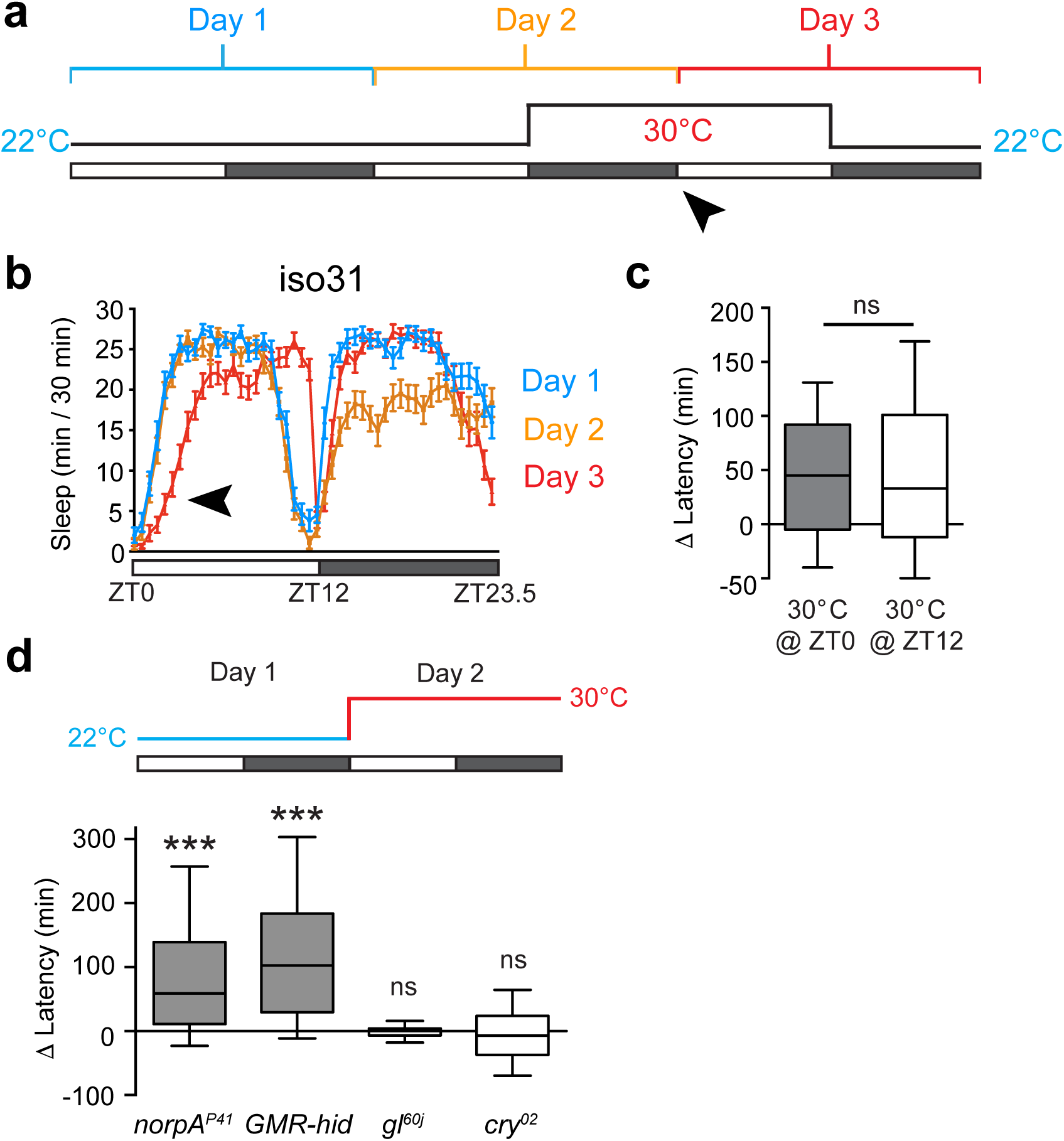
PMW is CRYPTOCHROME- and GLASS-dependent. (**a**) Three-day temperature-shift paradigm to test whether PMW is an acute avoidance response. Ambient temperature is raised from 22°C to 30°C at ZT12 on Day 2 for 24 h. Sleep latency was subsequently measured on Day 3 (black arrowhead). (**b**) Average sleep patterns of control adult male flies during the above temperature-shift paradigm. Subsequent days are juxtaposed to allow direct comparison. Day 1: blue, Day 2: orange, Day 3: red. Black arrowhead indicates PMW during Day 3. n = 43. (**c**) Comparison of the PMW when ambient temperature is increased at either ZT0 on the experimental day, or at ZT12 – the beginning of the previous night. ns – p > 0.05, Mann-Whitney U-test. ZT0: n = 79 (data also presented in Figure 1); ZT12: n = 45. (**d**) PMW in light-pathway mutants. Statistical comparison: Wilcoxon signed rank test compared to a theoretical median of zero. *norpA*^*P41*^: n = 31, GMR-*hid*: n = 38, *gl*^*60j*^: n = 44, *cry*^*02*^: n = 58.

### PMW is GLASS- and CRYPTOCHROME-dependent

Since PMW occurs shortly after lights-on, we tested whether PMW could be modified by mutations that impact light-sensing pathways, the circadian clock, or both (Fig. 2d). We examined PMW in three photoreceptor mutants where signaling through the compound eye is abolished. *norpA*^P41^ is a loss of function allele in the phospholipase C-β-encoding gene *norpA*, a critical component in the canonical light transduction pathway^24,25^; GMR-*hid* flies express the pro-apoptotic gene *hid* in all photoreceptor cells^26^; and *gl*^60j^ mutants are developmentally blind due to loss of GLASS, a transcription factor required for photoreceptor cell development^27^. Surprisingly, PMW was still observed in *norpA*^*P41*^ and GMR-*hid* males, yet was suppressed in *gl*^60j^ homozygotes (Fig. 2d). GLASS is also required for the development of a subset of clock cells termed DN1p neurons^26,28^, providing a possible explanation for this discrepancy (see below). Interestingly, we also found that PMW was suppressed by loss of CRYPTOCHROME (CRY), a circadian blue-light photoreceptor^29^ (Fig. 2d; see Discussion). Finally, to test for a direct role of the clock in gating PMW, we generated a new *timeless* (*tim*) null allele by replacing the *tim* coding sequence with a mini-*white*^*+*^ reporter gene using homologous recombination (*tim*^*ko*^; see Materials and Methods). TIM is an essential component of the negative arm of the circadian transcription-translation feedback loop^30^. We confirmed that clock-driven morning and evening anticipation are lost in *tim*^ko^ homozygote males (Fig. S2), and that *tim*^ko^ homozygotes are arrhythmic in constant-dark conditions (data not shown). PMW was reduced in *tim*^*ko*^ homozygotes, but not fully suppressed (Fig. S2). These results suggest that light- and temperature information collectively drive PMW, with the light-sensing pathway involving the CRY photoreceptor. Furthermore, the circadian clock may play a modulatory role in this process.

### PMW requires the TrpA1 thermo-receptor

What molecular pathways signal sleep-relevant temperature information to regulate siesta onset? The *Drosophila* genome encodes several thermo-sensory proteins^31^. Of these, TrpA1, a cation-conducting channel active at 30°C, is required for temperature-induced changes in the phase of morning and evening anticipation^23^. We therefore tested whether TrpA1 also impacted PMW. Indeed, a loss of function mutation in TrpA1 (*TrpA1*^1^) suppressed PMW, while PMW was still robustly observed in a paired genetic control (Fig. 3a-c). These results were confirmed using a previously validated *TrpA1-*RNAi^32^ transgene to knock down TrpA1 expression throughout the *Drosophila* nervous system using the pan-neuronal driver *elav*-GAL4 (Fig. S3). In the TrpA1 knockdown background, PMW was suppressed (Fig. S3), similarly to *TrpA1*^1^ homozygotes (Fig. 3a-c). In contrast, null or strongly hypomorphic mutations affecting the Gr28b^33^ and Pyrexia^34,35^ thermo-receptors did not suppress PMW (Fig. S3). From the above data, we conclude that TrpA1 is the critical thermo-sensor that mediates PMW.

**Figure 3:**
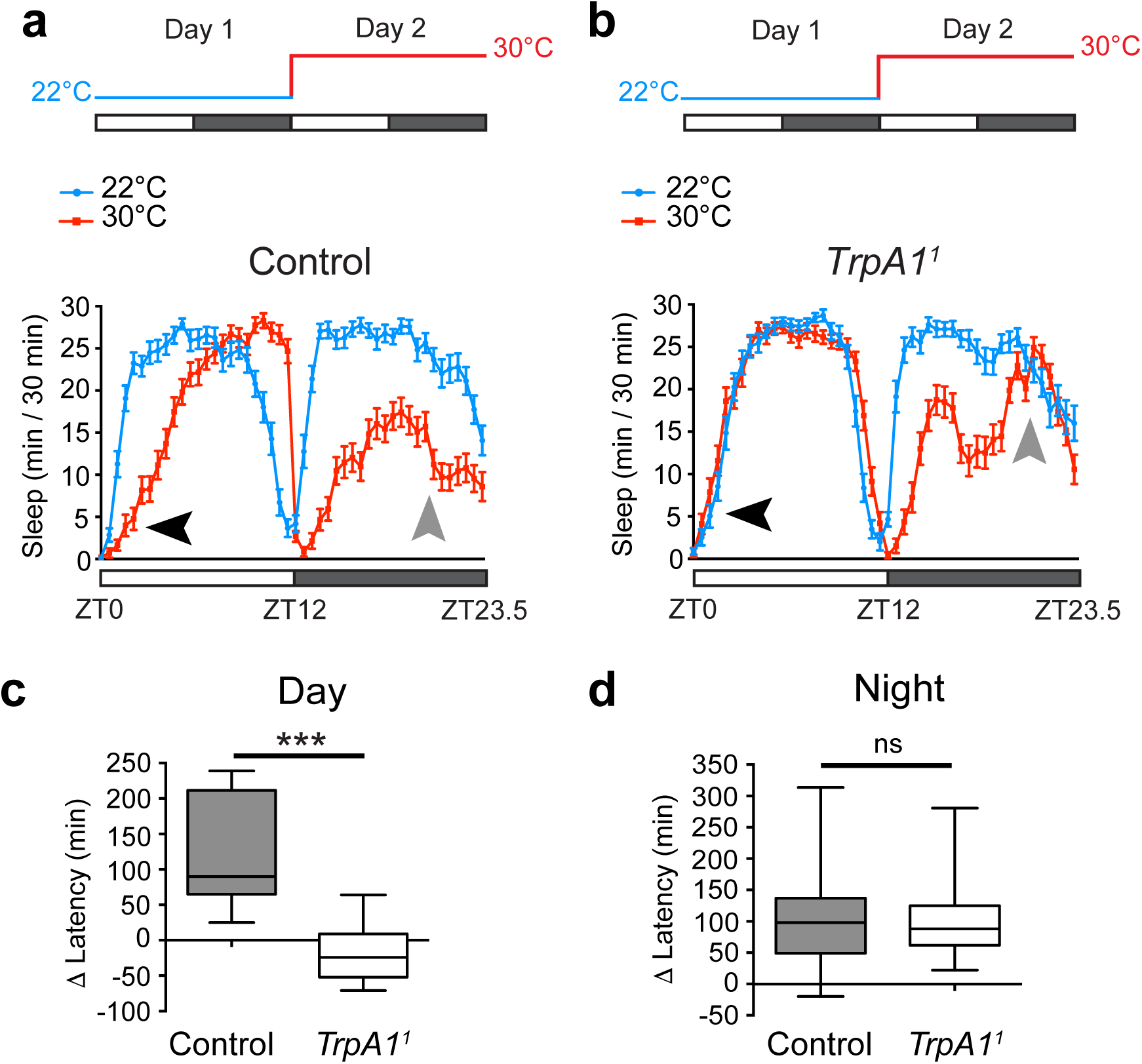
The TrpA1 thermo-sensor is required for PMW. (**a-b**) Average sleep patterns of adult male control or *TrpA1*^1^ homozygotes shifted from 22°C to 30°C at ZT0. Temperature-shift paradigms are indicated above. Black arrowheads: presence/absence of PMW. Grey arrowheads: presence/absence of enhanced wakefulness prior to lights-on during a warm night. (**c-d**) Comparison of change in latency to the first sleep episode between control and *TrpA1*^1^ homozygote males during the day (**c**) and night (**d**) following a shift from 22°C to 30°C. Statistical comparison: Mann-Whitney U-test. Control: n = 43; *TrpA1*^*1*^: n = 62.

As shown previously^23^, loss of TrpA1 also suppressed temperature-induced changes in morning and evening anticipation (Fig. 3a, b), but had no effect on temperature-induced delay of sleep onset during the night (Fig. 3d). Furthermore, sleep loss at 30°C was still strongly observed in the early-middle of the night in *TrpA1*^1^ mutants (Fig. 3b). Thus, temperature-dependent modulation of day sleep, but not night sleep, appears strongly dependent on TrpA1.

### Two populations of TrpA1-expressing neurons are necessary for PMW

We next sought to identify subpopulations of TrpA1-expressing neurons that transduce thermo-sensory information to drive PMW. Recent work has shown that a group of TrpA1-expressing neurons defined by the *TrpA1*[SH]-GAL4 driver modulates the phase of morning anticipation in response to temperature changes^23^. To test if these neurons also play a role in PMW, we used *TrpA1*-RNAi to knockdown TrpA1 in *TrpA1*[SH]-neurons. Indeed, PMW was reduced when TrpA1 expression was inhibited in *TrpA1*[SH]-neurons (Fig. 4a, b, d). To test whether additional TrpA1-expressing neurons also influenced PMW we screened several driver lines shown, or predicted, to label TrpA1-positive neurons. From this mini-screen, we found that expression of *TrpA1*-RNAi using *pickpocket*-GAL4 (*ppk*-GAL4), also robustly suppressed PMW (Fig. 4a, c, d). We further found that acute inhibition of *TrpA1*[SH]- and *ppk-*neuron output using temperature-sensitive dominant-negative *shibire* (UAS-*shi*^ts^), which blocks endocytosis of synaptic vesicles at 30°C but not 22°C^36^, also suppressed PMW (Fig. 4e). However, neither TrpA1 knockdown nor inhibition of synaptic transmission in *TrpA1*[SH]- and *ppk-*neurons suppressed temperature-induced delays in nighttime sleep onset (Fig. S4). Thus, TrpA1 expression in, and neurotransmitter release from, *TrpA1*[SH]- and *ppk-*neurons are required for PMW, and these circuits primarily impact daytime, as opposed to nighttime, sleep onset.

**Figure 4:**
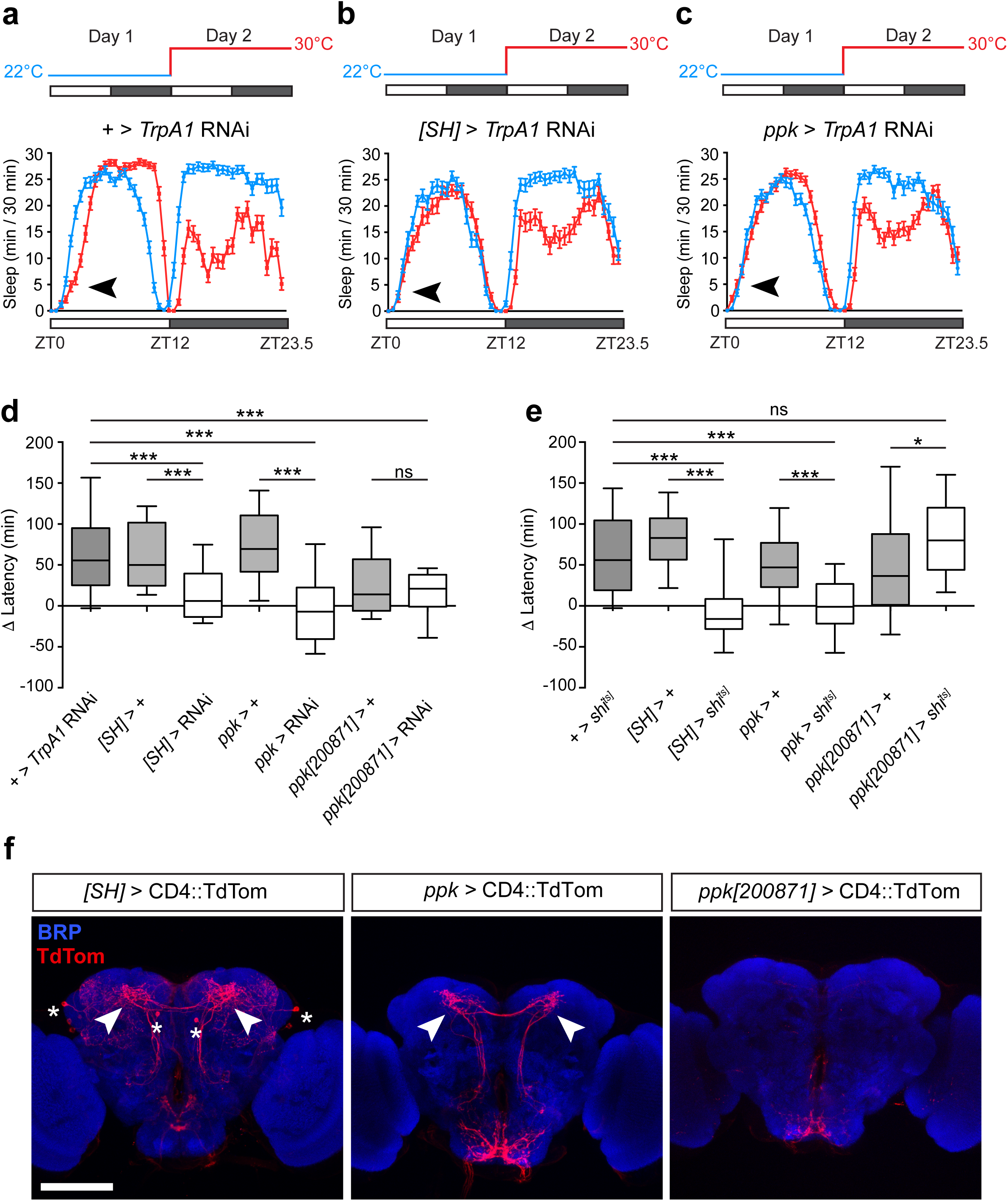
*TrpA1*-expressing *TrpA1*[SH]- and *ppk*-neurons are necessary for PMW. (**a-c**) Average sleep patterns of adult males containing a UAS-*TrpA1* RNAi transgene but lacking a *promoter*-GAL4 driver (**a**), or expressing UAS-*TrpA1* RNAi under control of the *TrpA1*[SH]- and *ppk*-GAL4 drivers (**b** and **c** respectively). Temperature-shift paradigms are indicated above. Black arrowheads: presence/absence of PMW. (**d**) Comparison of PMW in males expressing UAS*-TrpA1* RNAi under GAL4 drivers and their corresponding controls. n = 32-120. Statistical comparison: Kruskal-Wallis test with Dunn’s post-hoc test. All controls are ***p < 0.0001. *[SH]* > RNAi: p = 0.0481; *ppk* > RNAi: p = 0.2420; *ppk[200871]* > RNAi: p = 0.0054, using Wilcoxon signed rank test compared to a theoretical median of zero. (**e**) Effect of acute inhibition of synaptic output (using UAS-*shi*^ts^) from *TrpA1*[SH]-, *ppk*-, and *ppk[200871]*-neurons on PMW; Kruskal-Wallis test with Dunn’s post-hoc test. n = 24-77. All controls except *ppk[200871]* > + (**p = 0.0008) are ***p < 0.0001. *[SH] > shi*^*ts*^: ns (p = 0.06); *ppk > shi*^*ts*^: ns (p = 0.89); *ppk[200871] > shi*^*ts*^: ***p < 0.0001, Wilcoxon signed rank test compared to a theoretical median of zero. (**f**) Expression patterns of *TrpA1*[SH]*-, ppk-* and *ppk[200871]*- GAL4 in the adult *Drosophila* brain. TdTom labeled with anti DsRed. Synaptic neuropil (BRP) is labeled using an anti-nc82 antibody. Stars: cell bodies labeled by *TrpA1*[SH]-GAL4. Arrows: projections to the dorsal posterior protocerebrum from subsets of *TrpA1*[SH]- and *ppk-*neurons. Similar projection were not observed in *ppk[200871]*-positive neurons. Scale bar: 100 μm.

What are the neuro-anatomical correlates of PMW suppression through blocking TrpA1-signaling? *ppk*-GAL4 is widely used for labeling TrpA1-expressing sensory class IV multi-dendritic (mdIV) neurons in the larval and adult body wall^37,38^, but is also expressed in additional neurons in the adult legs, wings and antennae (data not shown). Interestingly, comparison of the projection patterns of *TrpA1*[SH]- and *ppk-*neurons in the adult brain suggested a potential commonality: both drivers encompass neurons that project to the dorsal posterior protocerebrum (DPP: Fig. 4f, arrows). As part of our mini-screen we also tested eight promoter fragments of the *ppk*-promoter fused to GAL4 (see Materials and Methods). Of these, the *ppk*[200871]-GAL4 driver labeled mdIV neurons on the adult body wall and exhibited a similar projection pattern to *ppk*-GAL4 in the suboesophageal ganglion (SOG) region of the brain (Fig. 4f and Fig. S4). However, projections to the DPP were absent in *ppk*[200871]-neurons, and both knockdown of TrpA1 and acute inhibition of synaptic output in *ppk*[200871]-neurons did not impact PMW (Fig. 4d, e). Thus, while we are yet to identify the precise cells within the *TrpA1*[SH]- and *ppk-*neuron populations that mediate PMW, the above results suggest that *TrpA1*[SH]- and non-mdIV *ppk-*positive neurons that project to the DPP may be critical mediators of PMW.

### Dorsal-projecting *TrpA1*[SH]- and *ppk*-neurons are distinct cell-types

Since both the *TrpA1*[SH]- and *ppk-*positive populations include neurons that send projections to the DPP and regulate PMW, we wondered whether the *TrpA1*[SH]- and *ppk-*GAL4 drivers label a common set of sensory neurons. *TrpA1*[SH]-GAL4 labels internal thermo-sensory AC neurons, whose axons project to the DPP from cell bodies located close to the antennal lobes^39^. In support of the above premise, we stochastically observed AC cell bodies when examining fluorescently labeled *TrpA1*[SH]- and *ppk-*neurons (Fig. S5). Thus, we used an intersectional strategy to provide more definitive evidence for common, or distinct, circuits labeled by *TrpA1*[SH]- and *ppk-*GAL4. We drove expression of the GAL4-inhibitory protein GAL80 under control of the *ppk*-promoter (*ppk*-GAL80^40^). In the presence of *ppk*-GAL80, we observed a strongly penetrant loss of *ppk*-GAL4 expression using CD4::TdTomato (CD4::TdTom) as a fluorescent reporter (Fig. 5a). We confirmed suppression of *ppk*-GAL4 by *ppk*-GAL80 at the behavioral level by driving UAS-*shi*^ts^ in *ppk*-GAL4/*ppk*-GAL80 males. In this background we still observed robust PMW (Fig. 4e), in contrast to the effect of driving UAS-*shi*^ts^ with *ppk*-GAL4 in the absence of *ppk*-GAL80 (Fig. 4e). Thus, *ppk*-GAL80 robustly suppresses *ppk*-GAL4 activity. Is the same true for *TrpA1*[SH]-GAL4? Unlike *ppk*-GAL4, *TrpA1*[SH]-GAL4-driven CD4::TdTom fluorescence was clearly observed in the presence of *ppk*-GAL80 (Fig. 5a), and expression of UAS-*shi*^ts^ in a *TrpA1*[SH]-GAL4/*ppk*-GAL80 background still inhibited PMW at 30°C (Fig. 5b). These data demonstrate that the critical *TrpA1*[SH]- and *ppk*-positive neurons required for PMW are distinct populations.

**Figure 5:**
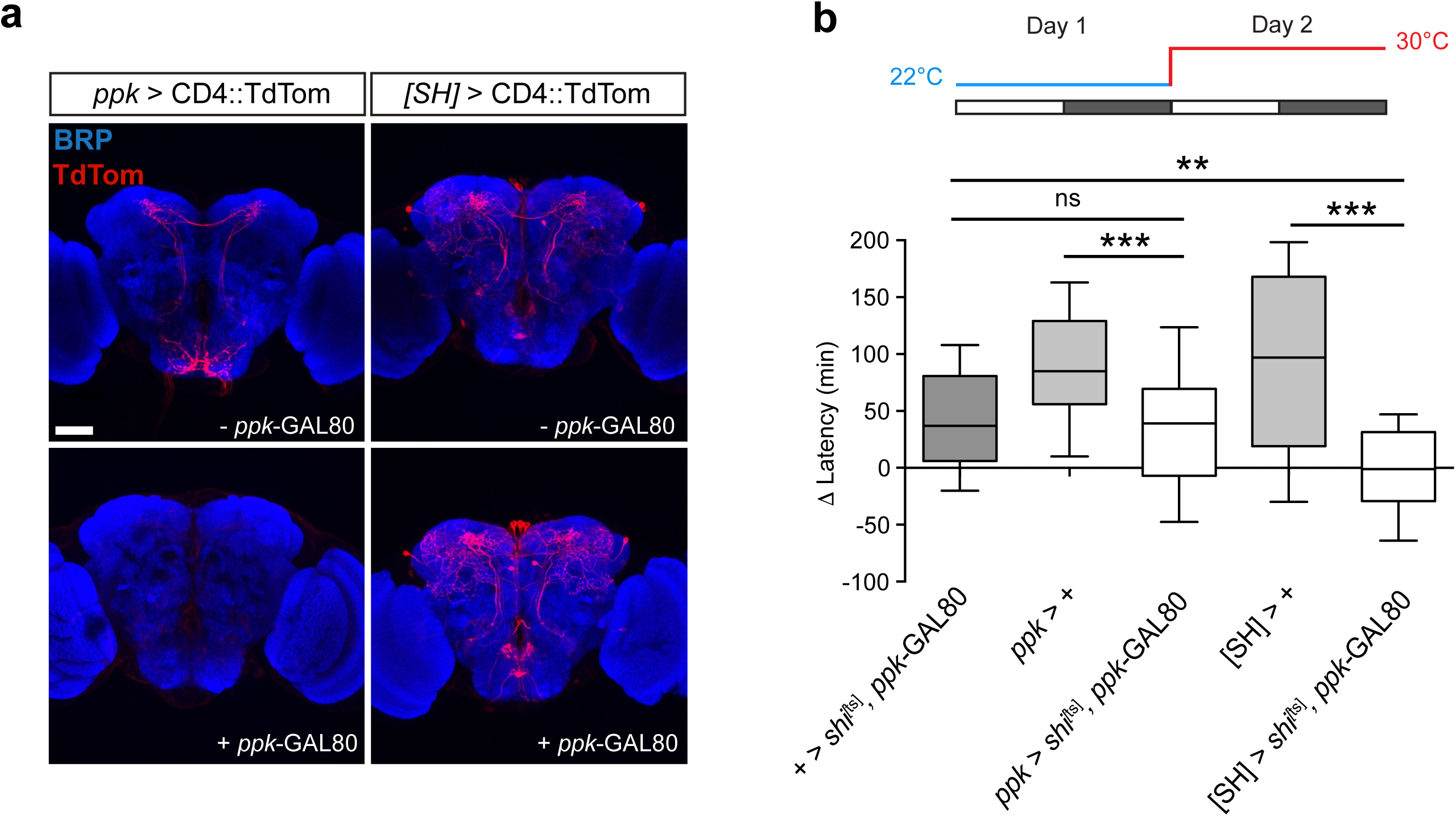
*TrpA1*[SH]- and *ppk*-neurons are distinct cellular populations. (**a**) Expression patterns of the *ppk-* and *TrpA1*[SH]-GAL4 drivers in adult male brains in the absence (top) or presence (bottom) of *ppk*-GAL80 - a GAL4-inhibitory protein under control of the *ppk*-promoter. Scale bar: 50 μm. (**b**) Effect of acute inhibition of synaptic output from *ppk*-and *TrpA1*[SH]-neurons on PMW using UAS-*shi*^ts^ in the presence of *ppk*-GAL80. n = 34-62, Kruskal-Wallis test with Dunn’s post-hoc test. All controls p < 0.0001; *ppk*>*shi^[ts]^, ppk*- GAL80: p=0.0005; *[SH]* > shi^*ts*^, *ppk*- GAL80: p = 0.91. Statistical comparison: Wilcoxon signed rank test compared to a theoretical median of zero.

### DN1p clock neurons are required for PMW and contact thermo-sensory neurons

Loss of the circadian photoreceptor CRY suppresses PMW (Fig. 2d). Therefore, we wondered whether subsets of clock neurons in the brain might regulate PMW. Since both *TrpA1*[SH]- and *ppk*-neurons send projections to the DPP, we focused on clock neurons with cell bodies and/or projections in this region. Within the network of clock neurons, CRY-positive s-LN_v_s, LN_d_s, and DN1p neurons send projections to the DPP^41^. DPP-projecting s-LN_v_s express the neuropeptide Pigment Dispersing Factor (PDF), which acts as a critical mediator of morning anticipation and rhythmicity in constant-dark conditions^42^, and both PDF-expressing s-LN_v_s, and DN1p neurons drive clock-dependent morning anticipation^43,44^. Interestingly, the output of DN1p neurons during the morning has also been shown to be temperature-dependent^44^, and the excitability of DN1p neurons peaks around dawn^45^, the time period in which PMW occurs (Fig. 1c). Furthermore, the development of DN1p neurons, but not s-LN_v_s, is GLASS-dependent^26,28^, and as shown above, loss of GLASS suppresses PMW (Fig 2d). Therefore, we tested for a direct role for DN1p neurons by acutely inhibiting DN1p synaptic output at 30°C by driving UAS-*shi*^ts^ with the driver *clk4.1M*-GAL4 (*4.1M*), which labels CRY-positive DN1p neurons in the adult brain^44^. When shifted to 30°C, inhibiting DN1p output suppressed PMW, whereas PMW was intact in control lines (Fig. 6a-d). In contrast, inhibiting DN1p output did not alter the delay in night sleep onset at 30°C (Fig. S6). Similar expression of UAS-*shi*^ts^ in the CRY-positive s-LN_v_s and LN_d_s using *mai179*-GAL4^43^ did not suppress PMW (Fig. S6). Since blocking classical neurotransmitter release does not inhibit PDF exocytosis^46^, we also tested for PMW in *pdf* null males (*pdf*^01^)^42^. In this background, PMW was present, albeit slightly reduced (Fig. S6). These results suggest that DN1p clock neurons are wake-promoting in the early morning at elevated temperatures and undertake a privileged role within the circadian CRY-positive network in regulating PMW.

**Figure 6:**
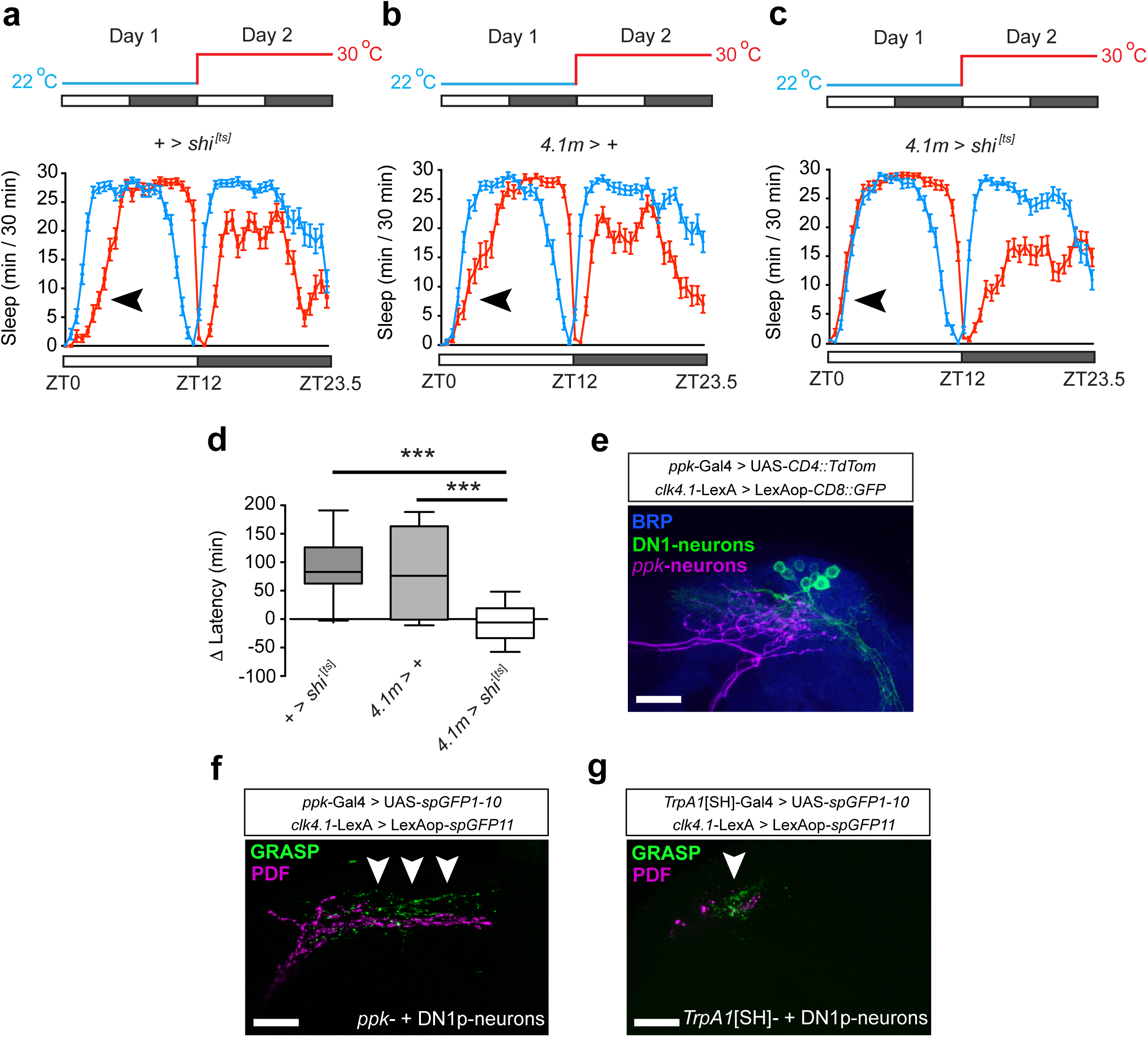
DN1p clock-neurons are necessary for PMW. (**a-c**) Average sleep patterns of adult males with synaptic output of DN1p neurons inhibited using UAS-*shi*^ts^ (*4.1M* > shi^ts^, **a**), and associated controls (**b-c**). Temperature-shift paradigms are indicated above. Black arrowheads: presence/absence of PMW. (**d**) Comparison of PMW in *4.1M* > *shi*^ts^ males and associated controls. n = 37-53, Kruskal-Wallis test with Dunn’s post-hoc test. All controls p < 0.0001; *4.1M* > *shi*^*ts*^: p = 0.24, using Wilcoxon signed rank test compared to a theoretical median of zero. (**e**) Co-localization of projections from *ppk*-neurons (magenta) and DN1p neurons (green) in the dorsal posterior protocerebrum of the adult *Drosophila* brain. BRP labeled with anti-nc82 antibody. (**f-g**) GRASP between DN1p neurons and *ppk*-neurons (**f**) or *TrpA1*[SH]-neurons (**g**). Arrows indicate regions of punctate GRASP signal. Scale bars: 20 μm.

DN1p cell bodies are located in the DPP, potentially in close proximity to projections from thermo-sensory *ppk*- and *TrpA1*[SH]-neurons. To confirm this, we used orthogonal binary systems to drive distinct fluorophores in both *ppk*- and DN1p neurons. Indeed, we observed an overlap between *ppk*- and DN1p-projections (Fig. 6e). To study potential connectivity between thermo-sensory and DN1p neurons, we used GRASP^47^ to test for physical interactions between *ppk*- and DN1p neurons, and *TrpA1*[SH]- and DN1p neurons. Using PDF antibody as a marker for the location of DN1p projections^48^, in both cases, expression of complementary split-GFP fragments in either *ppk*- and DN1p-neurons (Fig. 6f), or *TrpA1*[SH]- and DN1p-neurons (Fig. 6g) resulted in GRASP fluorescence. In contrast, we only observed minimal GRASP fluorescence between *ppk*- and PDF-neurons (Fig. S6). Collectively, these results suggest that DN1p-neurons receive dual input from two distinct populations of thermo-sensory neurons to drive temperature-induced increases in morning arousal.

## Discussion

How plasticity of distinct sleep periods is regulated at the molecular and circuit levels is unclear. Here we show that the TrpA1 thermo-sensor imparts temperature-sensitivity to siesta sleep in *Drosophila*, but modulates night sleep to a more subtle degree. Furthermore, we define a novel circuit linking thermo-sensory cells to clock neurons that, in turn, delay sleep onset in response to elevated temperatures.

Modulation of *Drosophila* sleep by temperature has recently been examined, but only up to an ambient temperature of 29°C^19,20^. At this temperature, siesta sleep was shown to increase relative to 25°C in both male and female flies^20^, and our results are consistent with this finding (Fig. 1f and Fig. S1). However, we find that in male flies, this effect is specific to the 27-29°C range. At higher temperatures that would nonetheless be common during summer months (≥ 30°C), siesta sleep is reduced. In particular, we observed a clear delay in siesta onset at ≥ 30°C that we term PMW (Fig. 1cd). The timing and magnitude of siesta sleep is sexually dimorphic^7^, with male flies initiating sleep earlier in the morning at 22°C when females are still active. While the causes of this sex-specific sleep pattern are still being elucidated, it is noticeable that females do not show PMW at 30°C. We suggest that the relative hyperactivity of females in the morning masks the effect of temperature on arousal, and later in the afternoon, circadian and/or homeostatic mechanisms act to initiate sleep, whether at mild or higher ambient temperatures. These results imply that arousal during the early morning is particularly sensitive to temperature increases. The circuits we have identified suggest an explanatory basis for this effect.

We found that TrpA1 acts in two distinct thermo-sensory subpopulations defined by the *TrpA1*[SH]- and *ppk*-GAL4 drivers to drive PMW, and that both DPP-projecting *TrpA1*[SH]- and *ppk*-neurons make physical contact with DN1p neurons that promote arousal in the early morning at 30°C (Figs. 4 and 6). When ectopically expressed, enhanced synaptic transmission induced by TrpA1 can be detected at 26°C and is further increased at 29°C^39^. We hypothesize that DN1p neurons receive weak excitatory drive from DPP-projecting *TrpA1*[SH]- and *ppk*-neurons, perhaps due to low TrpA1 expression or intrinsic excitability in each cell-type. In this model, excitatory drive scales with temperature^39^, and simultaneous input from *TrpA1*[SH]- and *ppk*-neurons, in combination with strong TrpA1-dependent activation of both circuits, is required to cause robust DN1p firing. This, in turn, prolongs arousal during morning periods.

Our model, combined with prior literature, suggests a mechanism for the relatively specific effect of TrpA1 signalling and DN1p activation on the onset of siesta, rather than night sleep. Under LD conditions, DN1p neurons promote morning anticipation, i.e increased locomotion before lights-on, and this output is reduced at low temperatures^44,49^. Recent work has also shown that thermo-genetic activation of CRY-positive DN1p neurons with a distinct driver (*R18H11*-GAL4^50^) induces a PMW-like phenotype (Fig. 4 of ^51^), further supporting a role for DN1p neurons in this process. Importantly, the intrinsic excitability of DN1p neurons is under circadian control, peaking between ZT20-ZT4 and reaching a minimum between ZT8-16 due to clock-dependent oscillations in the resting membrane potential^45^ (RMP). In parallel, PDF signalling from s-LN_v_s further enhances DN1p excitability in the late night/early morning^52,53^. Thus, DN1p neurons are ‘primed’ to receive excitatory input from thermo-sensory neurons during the early morning.

Consistent with clock- and PDF-dependent increases in DN1p excitability during the morning, we observed that loss of both the clock protein TIM, and PDF, reduce the magnitude of PMW (Fig. S2 and S6). We further demonstrate a role for the blue-light photoreceptor CRY as an essential molecular regulator of PMW (Fig. 2d). CRY is expressed in s-LN_v_, LN_d_ and DN1p clock neurons^41,54^, and *cry* transcription is clock-controlled^55^. Inhibiting neurotransmitter or neuropeptide release from the s-LN_v_s and LN_d_s does not phenocopy the effect of loss of CRY on PMW (Fig. 2 and Fig S6). In contrast, inhibiting DN1p output fully suppresses PMW (Fig. 6), as observed in *cry* null males (Fig. 2). Thus, the most parsimonious hypothesis is that CRY is acting in DN1p neurons.

How might CRY influence PMW, and since *cry* transcription is under clock control, why is PMW not fully suppressed in *tim* mutants? While CRY undertakes several roles in the *Drosophila* nervous system^29,56-59^, recent work has shown that CRY additionally mediates acute light-dependent increases in clock cell excitability via interaction with the potassium b-subunit Hyperkinetic^60,61^. CRY stability is light-dependent, and thus CRY protein levels increase during the night^55^. We hypothesize that in the early morning, strongly-expressed CRY confers light-dependent excitation to DN1p neurons, enhancing the effect of excitatory drive from TrpA1-expressing neurons. Loss of the negative feedback loop of the circadian clock results in constant low-level transcription of *cry*^55^. However, CRY protein may still accumulate during the night and promote PMW in the early morning. This may explain why loss of TIM reduces, but does not fully suppress, PMW (Fig. S2). Further experiments are required to test the predictions outlined above, and to identify the critical TrpA1-expressing cells that contact DN1p neurons.

In summary, we propose that DN1p neurons integrate both TrpA1-dependent temperature- and CRY-dependent light-information with clock-driven changes in intrinsic excitability to time sleep onset during the early day. In the wild, signalling from a wide array of sensory modalities must be computed in parallel to match sleep onset with environmental conditions. Our work provides a framework to unravel how multi-sensory processing in the *Drosophila* nervous system facilitates dynamic control of sleep timing.

## Materials and Methods

### Fly Strains and Husbandry

Fly strains and crosses were reared on standard yeast-containing fly flood at constant temperature 25°C in 12h: 12h Light-Dark cycles (LD).The *ppk*-Gal4 (BL32078), UAS-*shi*^[ts]^ (BL44222), UAS-*TrpA1* (BL26263), *TrpA1*^1^ (BL26504), and *gl*^*60j*^ (BL509) were obtained from Bloomington stock center. The *TrpA1*[SH]-Gal4, *pyx*^3^, *cry*^02^, GMR-*hid*, *norpA*^*P41*^, *pdf*^*01*^, *Clk*4.1M-LexA, UAS-*spGFP1-10* and LexAop-s*pGFP11* fly lines were described previously^24,26,34,39,42,47,48,62^. Apart from *gl*^*60j*^, *norpA*^*P41*^ and *pdf*^*01*^, all *Drosophila* lines used for sleep studies were outcrossed at least 5 times into an isogenic (iso31) background. This stock also served as a wild type control line. For the *gl*^*60j*^ and *pdf*^01^ stocks, chromosomes X and II were replaced with iso31 counterparts, while the original chromosome III (containing the *glass* or the *pdf* allele) was retained. The Vienna Tile (VT) lines were obtained from the Vienna *Drosophila* Resource Centre (VDRC). VDRC ID: 200748, 200871, 200905, 201215, 207296, 213670, 205646, 200782. VT insertions were also outcrossed 5 times into an iso31 background. 200871 and 207296 exhibit the most similar projection patterns in brain compared to *ppk*-Gal4 (see Fig. 4 and S4 for description of 200871). However, 207296 presents, in addition to the *ppk*-like SOG projections (and mdIV expression), expression in Giant Commisural Interneurons.

### Generation of the *timeless* knockout allele

The *timeless* knock-out fly line was generated using homologous recombination^63^. Homologous regions flanking the target gene locus were amplified via PCR, using *w*^1118^ genomic DNA as a template and primers containing appropriate restriction sites for cloning into the pGX-*attP* vector ^64^. Primer sequences were as follows. (5’ homologous region (4124 bp): 5’-ATAGCGGCCGCGAAGATTGTATACTCTAGAAG-3’/5’-CGATGGTACCATACCCTAATCGAAGTTGGTT-3’, *NotI/KpnI*. 3’ homologous region (3666 bp): 5’-ACGTACTAGTAACTGTGCAGGATATACGAATC-3’/5’-ATCGCTCGAGGGTCAAGATCTATTGGGAGTT-3’, *SpeI/XhoI*). pGX-*attP* was incorporated into the genome of *w*^1118^ flies via *P*-element insertion (BestGene Inc., and mapped to the second chromosome (BestGene Inc., California, USA). Ends-out targeting was performed as described previously^63-66^. Successful targeting events generated a deletion of ∼ 17 kb, including putative 5’ promoter and 3’ UTR sequences. Fly lines required to initiate homologous recombination were obtained from the Bloomington Drosophila Stock Center (BL#25679, BL#26258, BL#1092). After screening for potential targeting events (using a mini-*white*^+^ reporter as a visual marker), knockout was verified via genomic, proteomic and behavioural investigations (data not shown and Fig. S2).

### Behavioral assays

Individual 2-4 day old males were loaded into glass tubes (Trikinetics) containing 2% agar and 4% sucrose. For all experiments shown in this manuscript, Trikinetics monitors were housed in temperature- and light-controlled incubators (LMS, UK). Light intensity was measured to be between 700-1000 Lux using an environmental monitor (Trikinetics). Locomotor activity was recorded in 1 min bins using the *Drosophila* Activity Monitoring (DAM) system (Trikinetics). During temperature-shift experiments, flies were left for 22°C prior to recording for 24 h. Activity counts were subsequently measured for 24 h at 22°C, then for 24 h at elevated temperature (27°C, 29-31°C). The time taken for ambient temperature to increase from 22°C to 30°C was approximately 25 min. Sleep was defined as a period of inactivity of at least 5 min^67^. A modified version of a previously described Microsoft Excel script^68^ was used to measure all sleep parameters detailed in this article. Siesta sleep onset is defined as the latency of the first sleep bout and siesta offset as the end of the last sleep episode. We note that, using incubators from LMS (UK) in the lab of J.E.C.J, consistent and significant delays in siesta onset were observed in response to increasing ambient temperature from 22°C to 30°C in all control lines. A delayed onset using Percival DR36VLC8 incubators (IA, USA) was also observed in the lab of K.K, although the magnitude of the effect was marginally smaller. Thus, at 30°C, differences in incubator model may contribute to slightly altered effect sizes. When shifting flies from 22°C to 31°C, substantial and highly significant delays in siesta onset were observed using both incubator models. Sleep graphs were generated using GraphPad Prism 6.

### Immunohistochemistry and confocal microscopy

Adult male *Drosophila* brains were immuno-stained as described previously^10^. Briefly, brains were fixed in 4% paraformaldehyde for 20 min at RT, and blocked in 5% goat serum for 1 h at RT. Primary antibodies used were as follows: rabbit anti-DsRed (Clontech) – 1:2000; mouse anti-PDF (Developmental Studies Hybridoma Bank, DSHB) – 1:2000; mouse anti-Bruchpilot (nc82, DSHB) – 1:200; chicken anti-GFP (Invitrogen) – 1:1000. Alexa-fluor secondary antibodies were (goat anti-rabbit 555, goat anti-chicken 488 and goat anti-mouse 647, Invitrogen) were used at 1:2000 except for labelling anti-BRP where goat anti-mouse 647 where a dilution of 1:500 was used. Confocal images were taken using an inverted Zeiss LSM 710.

### Statistical analysis

Since many of the datasets derived from sleep experiments exhibited a non-normal distribution, the following statistical tests were used. Firstly, for binary analysis of whether temperature increases did or did not alter a given sleep parameter, Wilcoxon signed rank test was used, with experimental medians compared to a theoretical median of zero. When simultaneously comparing multiple genotypes, Kruskal-Wallis tests were used, followed by Dunn’s post-hoc tests. All statistical analysis was performed using GraphPad Prism 6.

## Acknowledgments

We thank Dr. Ko-Fan Chen for generating the Excel script used to measure sleep onset. We thank Drs. Ralf Stanewsky, Nirao Shah, Yuh Nung Jan, Francois Rouyer and the Bloomington Stock Center for providing fly stocks. Dr. Ralf Stanewsky and Dr. Ko-Fan Chen gave helpful comments on the manuscript, and Abhishek Chatterjee gave helpful suggestions during the research process. Dr. Ralf Stanewsky also gave infrastructure support to J.E.C.J when initially constructing his lab. Huihui Pan provided technical support, and Alexandra Kenny performed pilot experiments. This work was supported by a grant from the National Institutes of Health (R01NS086887) to K.K and a UCL start-up grant to J.E.C.J.

### Author contributions

Conceptualization, A.L and J.E.C.J; Methodology, A.L and J.E.C.J; Formal Analysis, A.L; Investigation, A.L and A.O-C; Resources, R.F and N.P; Writing – Original Draft, J.E.C.J and A.L; Writing – Review & Editing; A.O-C, N.P and K.K; Visualization, A.L, A.O-C, K.K and J.E.C.J; Funding Acquisition, K.K and J.E.C.J; Supervision, K.K and J.E.C.J.

### Competing financial interests

The authors declare no competing financial interest.

## Reference

1 Borbély, A. A. A two process model of sleep regulation. Hum Neurobiol 1, 195–204. (1982).

2 Sehgal, A. & Mignot, E. Genetics of sleep and sleep disorders. Cell 146, 194–207 doi:10.1016/j.cell.2011.07.004 (2011).

3 Bushey, D., Tononi, G. & Cirelli, C. Sleep and synaptic homeostasis: structural evidence in *Drosophila*. Science 332, 1576–1581 doi:10.1126/science.1202839 (2011).

4 Gilestro, G. F., Tononi, G. & Cirelli, C. Widespread changes in synaptic markers as a function of sleep and wakefulness in *Drosophila*. Science 324, 109–112 doi:10.1126/science.1166673 (2009).

5 Kayser, M. S., Yue, Z. & Sehgal, A. A critical period of sleep for development of courtship circuitry and behavior in *Drosophila*. Science 344, 269–274 doi:10.1126/science.1250553 (2014).

6 Xie, L. et al. Sleep drives metabolite clearance from the adult brain. Science 342, 373–377 doi:10.1126/science.1241224 (2013).

7 Cirelli, C. et al. Reduced sleep in *Drosophila* Shaker mutants. Nature 434, 1087–1092 doi:10.1038/nature03486 (2005).

8 Koh, K. et al. Identification of SLEEPLESS, a sleep-promoting factor. Science 321, 372–376 doi:10.1126/science.1155942 (2008).

9 Liu, Q., Liu, S., Kodama, L., Driscoll, M. R. & Wu, M. N. Two dopaminergic neurons signal to the dorsal fan-shaped body to promote wakefulness in *Drosophila*. Curr Biol 22, 2114–2123 doi:10.1016/j.cub.2012.09.008 (2012).

10 Liu, S. et al. WIDE AWAKE mediates the circadian timing of sleep onset. Neuron 82, 151–166 doi:10.1016/j.neuron.2014.01.040 (2014).

11 Liu, S., Liu, Q., Tabuchi, M. & Wu, M. N. Sleep Drive Is Encoded by Neural Plastic Changes in a Dedicated Circuit. Cell 165, 1347–1360 doi:10.1016/j.cell.2016.04.013 (2016).

12 Ueno, T. et al. Identification of a dopamine pathway that regulates sleep and arousal in *Drosophila*. Nat Neurosci 15, 1516–1523, doi:10.1038/nn.3238 (2012).

13 Donlea, J. M., Pimentel, D. & Miesenbock, G. Neuronal machinery of sleep homeostasis in *Drosophila*. Neuron 81, 860–872, doi:10.1016/j.neuron.2013.12.013 (2014).

14 Pimentel, D. et al. Operation of a homeostatic sleep switch. Nature 536, 333–337, doi:10.1038/nature19055 (2016).

15 Berry, J. A., Cervantes-Sandoval, I., Chakraborty, M. & Davis, R. L. Sleep Facilitates Memory by Blocking Dopamine Neuron-Mediated Forgetting. Cell 161, 1656–1667, doi:10.1016/j.cell.2015.05.027 (2015).

16 Fowler, M. A. & Montell, C. *Drosophila* TRP channels and animal behavior. Life Sci 92, 394–403, doi:10.1016/j.lfs.2012.07.029 (2013).

17 Garrity, P. A., Goodman, M. B., Samuel, A. D. & Sengupta, P. Running hot and cold: behavioral strategies, neural circuits, and the molecular machinery for thermotaxis in *C. elegans* and *Drosophila*. Genes Dev 24, 2365–2382, doi:10.1101/gad.1953710 (2010).

18 Ishimoto, H., Lark, A. R. & Kitamoto, T. Factors that differentially affect daytime and nighttime sleep in Drosophila melanogaster. Front Neurol 3, 24 doi:10.3389/fneur.2012.00024 (2012).

19 Cao, W. & Edery, I. A novel pathway for sensory-mediated arousal involves splicing of an intron in the period clock gene. Sleep 38, 41 doi:10.5665/sleep.4322 (2015).

20 Parisky, K. M., Rivera, J. L. A., Donelson, N. C., Kotecha, S. & Griffith, L. C. Reorganization of Sleep by Temperature in *Drosophila* Requires Light, the Homeostat, and the Circadian Clock. Curr Biol 26, 882–892 doi:10.1016/j.cub.2016.02.011 (2016).

21 Pfeiffenberger, C., Lear, B. C., Keegan, K. P. & Allada, R. Processing sleep data created with the *Drosophila* Activity Monitoring (DAM) System. Cold Spring Harb Protoc 2010, pdb.prot5520, doi:10.1101/pdb.prot5520 (2010).

22 Majercak, J., Sidote, D., Hardin, P. E. & Edery, I. How a circadian clock adapts to seasonal decreases in temperature and day length. Neuron 24, 219–230 (1999).

23 Das, A., Holmes, T. C. & Sheeba, V. dTRPA1 in Non-circadian Neurons Modulates Temperature-dependent Rhythmic Activity in *Drosophila* melanogaster. J Biol Rhythms 31, 272–288 doi:10.1177/0748730415627037 (2016).

24 Szular, J. et al. Rhodopsin 5–and Rhodopsin 6–Mediated Clock. Synchronization in *Drosophila melanogaster* Is Independent of Retinal Phospholipase C-β Signaling. J Biol Rhythms 27, 25–36 doi:10.1177/0748730411431673 (2012).

25 Montell, C. *Drosophila* visual transduction. Trends Neurosci 35, 356–363 doi:10.1016/j.tins.2012.03.004 (2012).

26 Klarsfeld, A. et al. Novel features of cryptochrome-mediated photoreception in the brain circadian clock of *Drosophila*. J Neurosci 24, 1468–1477 doi:10.1523/JNEUROSCI.3661-03.2004 (2004).

27 Moses, K., Ellis, M. C. & Rubin, G. M. The glass gene encodes a zinc: finger protein required by *Drosophila* photoreceptor cells. Nature 340, 531–536 doi:10.1038/340531a0 (1989).

28 Helfrich-Förster, C., Winter, C., Hofbauer, A., Hall, J. C. & Stanewsky, R. The circadian clock of fruit flies is blind after elimination of all known photoreceptors. Neuron 30, 249–261 (2001).

29 Stanewsky, R. et al. The cryb mutation identifies cryptochrome as a circadian photoreceptor in *Drosophila*. Cell 95, 681–692 (1998).

30 Sehgal, A., Price, J. L., Man, B. & Young, M. W. Loss of circadian behavioral rhythms and per RNA oscillations in the *Drosophila* mutant timeless. Science 263, 1603–1605 (1994).

31 Barbagallo, B. & Garrity, P. A. Temperature sensation in *Drosophila*. Curr Opin Neurobiol 34, 8–13, doi:10.1016/j.conb.2015.01.002 (2015).

32 Green, E. W. et al. *Drosophila* circadian rhythms in seminatural environments: summer afternoon component is not an artifact and requires TrpA1 channels. Proc Natl Acad Sci 112, 8702–8707 doi:10.1073/pnas.1506093112 (2015).

33 Ni, L. et al. A gustatory receptor paralogue controls rapid warmth avoidance in *Drosophila*. Nature 500, 580–584 doi:10.1038/nature12390 (2013).

34 Wolfgang, W., Simoni, A., Gentile, C. & Stanewsky, R. The Pyrexia transient receptor potential channel mediates circadian clock synchronization to low temperature cycles in *Drosophila melanogaster*. Proc R Soc Lond B Bio Sci 280, 20130959 doi:10.1098/rspb.2013.0959 (2013).

35 Lee, Y. et al. Pyrexia is a new thermal transient receptor potential channel endowing tolerance to high temperatures in *Drosophila melanogaster*. Nat Genet 37, 305–310 doi:10.1038/ng1513 (2005).

36 Kitamoto, T. Conditional modification of behavior in *Drosophila* by targeted expression of a temperature-sensitive shibire allele in defined neurons. J Neurobiol 47, 81–92 (2001).

37 Grueber, W. B. et al. Projections of *Drosophila* multidendritic neurons in the central nervous system: links with peripheral dendrite morphology. Development 134, 55–64 doi:10.1242/dev.02666 (2007).

38 Xiang, Y. et al. Light-avoidance-mediating photoreceptors tile the *Drosophila* larval body wall. Nature 468, 921–926 doi:10.1038/nature09576 (2010).

39 Hamada, F. N. et al. An internal thermal sensor controlling temperature preference in *Drosophila*. Nature 454, 217–220 doi:10.1038/nature07001 (2008).

40 Yang, C.-h. et al. Control of the postmating behavioral switch in *Drosophila* females by internal sensory neurons. Neuron 61, 519–526 doi:10.1016/j.neuron.2008.12.021 (2009).

41 Yoshii, T., Todo, T., Wülbeck, C., Stanewsky, R. & Helfrich-Förster, C. Cryptochrome is present in the compound eyes and a subset of *Drosophila's* clock neurons. J Comp Neurol 508, 952–966 doi:10.1002/cne.21702 (2008).

42 Renn, S. C., Park, J. H., Rosbash, M., Hall, J. C. & Taghert, P. H. A pdf neuropeptide gene mutation and ablation of PDF neurons each cause severe abnormalities of behavioral circadian rhythms in *Drosophila*. Cell 99, 791–802 (1999).

43 Grima, B., Chélot, E., Xia, R. & Rouyer, F. Morning and evening peaks of activity rely on different clock neurons of the *Drosophila* brain. Nature 431, 869–873 doi:10.1038/nature02935 (2004).

44 Zhang, Y., Liu, Y., Bilodeau-Wentworth, D., Hardin, P. E. & Emery, P. Light and temperature control the contribution of specific DN1 neurons to *Drosophila* circadian behavior. Curr Biol 20, 600–605 doi10.1016/j.cub.2010.02.044 (2010).

45 Flourakis, M. et al. A conserved bicycle model for circadian clock control of membrane excitability. Cell 162, 836–848 doi:10.1016/j.cell.2015.07.036 (2015).

46 Johard, H. A. et al. Peptidergic clock neurons in *Drosophila*: ion transport peptide and short neuropeptide F in subsets of dorsal and ventral lateral neurons. J Comp Neurol 516, 59–73 doi:10.1002/cne.22099 (2009).

47 Feinberg, E. H. et al. GFP Reconstitution Across Synaptic Partners (GRASP) defines cell contacts and synapses in living nervous systems. Neuron 57, 353–363 doi:10.1016/j.neuron.2007.11.030 (2008).

48 Cavanaugh, D. J. et al. Identification of a circadian output circuit for rest: activity rhythms in *Drosophila*. Cell 157, 689–701 doi:10.1016/j.cell.2014.02.024 (2014).

49 Zhang, L. et al. DN1(p) circadian neurons coordinate acute light and PDF inputs to produce robust daily behavior in *Drosophila*. Curr Biol 20, 591–599 doi:10.1016/j.cub.2010.02.056 (2010).

50 Guo, F. et al. Circadian neuron feedback controls the *Drosophila* sleep–activity profile. Nature 536, 292–297 (2016). doi:10.1038/nature19097

51 Kunst, M. et al. Calcitonin gene-related peptide neurons mediate sleep-specific circadian output in *Drosophila*. Curr Biol 24, 2652–2664 doi:10.1016/j.cub.2014.09.077 (2014).

52 Seluzicki, A. et al. Dual PDF signaling pathways reset clocks via TIMELESS and acutely excite target neurons to control circadian behavior. PLoS Biol 12, e1001810, doi:10.1371/journal.pbio.1001810 (2014).

53 Park, J. H. et al. Differential regulation of circadian pacemaker output by separate clock genes in *Drosophila*. Proc Natl Acad Sci 97, 3608–3613, doi:10.1073/pnas.070036197 (2000).

54 Benito, J., Houl, J. H., Roman, G. W. & Hardin, P. E. The blue-light photoreceptor CRYPTOCHROME is expressed in a subset of circadian oscillator neurons in the *Drosophila* CNS. J Biol Rhythms 23, 296–307 doi:10.1177/0748730408318588 (2008).

55 Emery, P., So, W. V., Kaneko, M., Hall, J. C. & Rosbash, M. CRY, a Drosophila clock and light-regulated cryptochrome, is a major contributor to circadian rhythm resetting and photosensitivity. Cell 95, 669–679 (1998).

56 Kumar, S., Chen, D. & Sehgal, A. Dopamine acts through Cryptochrome to promote acute arousal in *Drosophila*. Genes Dev 26, 1224–1234 doi:10.1101/gad.186338.111 (2012).

57 Ceriani, M. F. et al. Light-dependent sequestration of TIMELESS by CRYPTOCHROME. Science 285, 553–556 (1999).

58 Gegear, R. J., Casselman, A., Waddell, S. & Reppert, S. M. Cryptochrome mediates light-dependent magnetosensitivity in *Drosophila*. Nature 454, 1014–1018 doi:10.1038/nature07183 (2008).

59 Mazzotta, G. et al. Fly cryptochrome and the visual system. Proc Natl Acad Sci 110, 6163–6168 doi:10.1073/pnas.1212317110 (2013).

60 Fogle, K. J. et al. CRYPTOCHROME-mediated phototransduction by modulation of the potassium ion channel beta-subunit redox sensor. Proc Natl Acad Sci 112, 2245–2250 doi:10.1073/pnas.1416586112 (2015).

61 Fogle, K. J., Parson, K. G., Dahm, N. A. & Holmes, T. C. CRYPTOCHROME is a blue-light sensor that regulates neuronal firing rate. Science 331, 1409–1413 doi:10.1126/science.1199702 (2011).

62 Dolezelova, E., Dolezel, D. & Hall, J. C. Rhythm defects caused by newly engineered null mutations in *Drosophila's* cryptochrome gene. Genetics 177, 329–345 doi:10.1534/genetics.107.076513 (2007).

63 Huang, J., Zhou, W., Dong, W., Watson, A. M. & Hong, Y. From the Cover: Directed, efficient, and versatile modifications of the *Drosophila* genome by genomic engineering. Proc Natl Acad Sci 106, 8284–8289 doi:10.1073/pnas.0900641106 (2009).

64 Huang, J., Zhou, W., Watson, A. M., Jan, Y. N. & Hong, Y. Efficient ends-out gene targeting in *Drosophila*. Genetics 180, 703–707 doi:10.1534/genetics.108.090563 (2008).

65 Gong, W. J. & Golic, K. G. Ends-out, or replacement, gene targeting in *Drosophila*. Proc Natl Acad Sci 100, 2556–2561 doi:10.1073/pnas.0535280100 (2003).

66 Maggert, K. A., Gong, W. J. & Golic, K. G. Methods for homologous recombination in *Drosophila*. Methods Mol Biol 420, 155–174 doi:10.1007/978-1-59745-583-1_9 (2008).

67 Shaw, P. J., Cirelli, C., Greenspan, R. J. & Tononi, G. Correlates of sleep and waking in *Drosophila melanogaster*. Science 287, 1834–1837 (2000).

68 Roessingh, S., Wolfgang, W. & Stanewsky, R. Loss of *Drosophila melanogaster* TRPA1 function affects “siesta” behavior but not synchronization to temperature cycles. J Biol Rhythms 30, 492–505 doi:10.1177/0748730415605633 (2015).

